# The Quest for Dynamic Consistency — A Comparison of OpenSim Tools for Residual Reduction in Simulations of Human Running

**DOI:** 10.1101/2023.08.31.555836

**Authors:** Aaron S. Fox

## Abstract

The use of synchronous kinematic and kinetic data in simulations of human running will typically lead to dynamic inconsistencies (i.e. residual forces and moments) being present. Minimising the residual forces and moments in such simulations is important to ensure plausible model outputs (e.g. joint moments, muscle forces) are obtained. A variety of approaches suitable for residual reduction are available in OpenSim, however a detailed comparison of these is yet to be conducted. This study compared a variety of OpenSim tools applicable for residual reduction in simulations of human running. A series of approaches (i.e. singular and iterative Residual Reduction Algorithm, *MocoTrack, AddBiomechanics*) designed to reduce residual forces and moments were examined using an existing dataset of 10 male participants running on a treadmill at 5.0 m·s^-1^ (*n* = 3 gait cycles per participant). The computational time, resultant residual forces and moments, and output joint kinematics and kinetics from each approach were compared. A computational cost to residual reduction trade-off was identified, where lower residual forces and moments were achieved using approaches that required longer computational times. All of the tested approaches regularly reduced residual forces below recommended thresholds, however only the *MocoTrack* approach could consistently achieve acceptable levels for residual moments. The *AddBiomechanics* and *MocoTrack* approaches produced variable lower and upper body kinematics, respectively, versus the remaining approaches; with minimal other qualitative differences were identified between joint kinematics from each approach. Joint kinetics were qualitatively similar between approaches, however *MocoTrack* generated much noisier joint kinetic signals. The *MocoTrack* approach was the most consistent and best performing approach for reducing residuals to near-zero levels, at the cost of longer computational times and potentially noisier joint kinetic signals. This study provides OpenSim users with evidence to inform decision-making at the residual reduction step of their modelling and simulation workflow when analysing human running.

## Introduction

Biomechanical data (i.e. kinematics and kinetics) are commonly collected and used alongside musculoskeletal models to understand human running performance or injury risk. Kinematic data is typically collected via marker-based optical motion capture systems, with kinetic data synchronously collected via in-ground force plates or instrumented treadmills. The independent measurement and associated error (i.e. noise) of kinematic and kinetic data in gait experiments leads to dynamic inconsistencies in modelled data [5]. Residual forces and moments at the ‘root’ segment (i.e. the segment connected to the ‘ground’ in a model — typically the pelvis) remain present to ensure dynamic consistency between the motion and external forces. The presence of dynamic inconsistencies can lead to implausible conclusions in simulation outputs (e.g. joint moments, muscle forces) given that not all of the forces are accounted for by realistic parts of the model. It is therefore common practice for biomechanists to employ strategies that minimise or eliminate residual forces and moments [5].

Adjusting model parameters (e.g. segment masses) and/or motion (e.g. joint kinematics) is a common approach for minimising residual forces and moments in gait simulations [5, 12]. OpenSim is a widely used modelling and simulation software, and offers the Residual Reduction Algorithm (RRA) as its main tool for minimising dynamic inconsistencies between modelled motions and external forces during gait [2]. RRA employs a forward dynamics simulation to adjust model kinematics and the mass centre of a selected body (typically the torso), while also providing recommendations for adjusting the mass of individual segments as a means to reduce residual forces and moments [1]. RRA can be effective in reducing residuals within recommended thresholds [5], however the process is dependent on selecting tracking weights for joint coordinates which may be difficult to objectively determine [9, 11]. Further, RRA has been employed as both a singular [4] and iterative [8] process — yet there are no specific recommendations on which of these approaches or how many iterations are optimal.

The expansion of OpenSim’s toolkit in recent years offers potential alternatives for generating dynamically consistent gait simulations. OpenSim Moco [3] provides an option to leverage direct collocation to achieve dynamic consistency — using the *MocoTrack* class to generate torque-driven simulations that track and adjust model kinematics, while minimising residual forces and moments. The recently released *AddBiomechanics* web application [12] aims to automate typical modelling processes (i.e. model scaling, inverse kinematics, inverse dynamics), and includes an optimisation step that updates model segment masses and joint kinematics to minimise dynamic inconsistencies in the final simulation results. A detailed comparison of available tools and their capacity to achieve dynamically consistent simulations of human running can provide researchers with information on which may be the most suitable approach(es). The purpose, nay quest, of this study was to compare the various OpenSim tools available for residual reduction in simulations of human running, with particular reference to the: (i) computational time; (ii) resultant residual forces and moments; and (iii) output joint kinematics and kinetics of each approach.

## Methods

### Dataset

This study used the human running dataset provided by Hamner and Delp [4], which includes ten male participants (age = 29 ± 5 y; height = 1.77 ± 0.04 m; mass = 70.9 ± 7.0 kg) running on a treadmill at three speeds (3.0 m·s^-1^, 4.0 m·s^-1^, 5.0 m·s^-1^). Only the data from participants running at 5.0 m·s^-1^ was used in the present study, given the fastest speed would likely include data with the highest forces and accelerations — and hence the greatest potential for residual forces and moments to be present in the experimental measurements. Data extracted from the original study [4] included the: (i) generic and participant-specific scaled full-body musculoskeletal models (12 segment, 29 degree-of-freedom musculoskeletal model); (ii) experimental marker and ground reaction force (GRF) data (i.e. .*trc* and .*mot* files); and (iii) full body joint coordinates from three gait cycles calculated via inverse kinematics (IK). All or parts of the extracted data were used in the subsequent residual reduction approaches tested.

### Data Analysis

The following sections outline the residual reduction approaches applied to the running dataset. All analyses were conducted in OpenSim 4.3 via Python 3.8 on a single CPU (11^th^ Gen Intel® Core™ i7-1185G7 processor; 16GB RAM with four cores), with the exception of the *AddBiomechanics* approach which were uploaded and processed in the web application [10]. The computational time (in minutes), average and peak residual forces (in N) and moments (in Nm) about the X (anterior-posterior), Y (medial-lateral) and Z (vertical) axes, and average whole-body joint kinematics and kinetics from the three gait cycles were extracted and descriptively compared across the residual reduction approaches. All data, analysis code and outputs can be accessed via the associated SimTK project page (https://simtk.org/projects/dynamic-quest).

### Residual Reduction Algorithm (RRA)

A single iteration of OpenSim’s RRA was implemented on the three gait cycles extracted for each participant using standardised practices. The inputs to the procedure were the scaled musculoskeletal model and experimental outputs (i.e. joint coordinates from IK and external GRFs), alongside the RRA settings files (i.e. joint coordinate tracking weights and model joint torque actuators) provided from the original study [4]. No adjustment to RRA settings on what were originally used by Hamner and Delp [4] were made to avoid introducing any further subjectivity to the process. A single iteration of the RRA was run on each gait cycle, providing the outputs of an adjusted musculoskeletal model (i.e. altered segment masses and torso mass centre) and joint coordinates. The residual forces and moments about the pelvis were determined from these outputs, alongside the computational time taken to complete the single RRA iteration — and averaged across each participants three gait cycles.

### Iterative Residual Reduction Algorithm (RRA3)

An iterative RRA approach [8] was also conducted, whereby three consecutive iterations of the RRA were conducted on the three gait cycles extracted for each participant. The same inputs outlined in the previous section were used in the first RRA iteration. Each further iteration of the RRA, however, used the adjusted musculoskeletal model and joint coordinates from the previous iteration alongside the original experimental GRFs and RRA settings files. The residual forces and moments about the pelvis from each gait cycle were determined from the final (i.e. third) RRA iteration musculoskeletal model and joint coordinate outputs, alongside the summed computational time taken to complete the three RRA iterations — and averaged across each participants three gait cycles.

### MocoTrack

Simulations of each participants three gait cycles were conducted using OpenSim’s *MocoTrack* class [3]. The optimal control simulations used a weighted (*w*) objective function with convergence and constraint tolerances of 1*e*^*−*2^ that minimised: (i) the joint coordinate tracking error with respect to the experimental IK input data (global *w* = 1); and (ii) the sum of squared torque actuator controls acting at each joint (global *w* = 1*e*^*−*3^), while also applying the experimental external GRFs. Settings which could be practically replicated across both the *MocoTrack* and RRA approaches were used in an attempt to ensure parity. This included using identical tracking weights for individual joint coordinates and optimal forces for torque actuators within the overall tracking and control goals, respectively. Replicating the time step of the RRA approach (i.e. 0.0001s) in the *MocoTrack* problem resulted in an extremely fine mesh interval which would have taken an impractically long duration to solve. The number of collocation nodes used in the *MocoTrack* problem was therefore determined using a mesh interval of 0.01s. While *MocoTrack* has an ability to include parameter optimisation which could be applied to segment masses of the musculoskeletal model, this was not considered in the present study. The residual forces and moments about the pelvis from each gait cycle were determined from the converged *MocoTrack* simulations, alongside the computational time taken for the problem to solve — and averaged across each participants three gait cycles.

### AddBiomechanics

*AddBiomechanics* [12] is an online application which provides automated processing of experimental marker and GRF data. It includes an initial model scaling and inverse kinematics step to produce joint coordinates of the input motion that minimise marker error. Following this, a second optimisation can be run alongside the inverse dynamics step which aims to refine body segment masses and joint coordinates to produce a more dynamically consistent motion. The outputs from this second optimisation are therefore of the most interest to the present study. The entire *AddBiomechanics* pipeline is outlined in Werling et al. [12], and hence the specific details are not provided here.

The input data for the *AddBiomechanics* approach differed to those previously outlined, where: (i) the unprocessed experimental marker and GRF data were used; and (ii) the entire running trial was used rather than being separated into gait cycles. On the latter point — the *AddBiomechanics* application suggests movement trials that include a large range of movement are optimal, and will subsequently prompt users with a warning when minimal frames of data (e.g. from a single gait cycle) are provided. The experimental marker and GRF data for each participant were uploaded to the *AddBiomechanics* application for processing. The option to use a custom musculoskeletal model and markerset were selected to ensure consistency with the previous approaches, while all other settings (e.g. the weight of residuals in the main optimisation) were kept as their default. The option to run an additional optimisation to try and drive residuals to exactly zero at the cost of more marker error was selected. However, the *AddBiomechanics* application notes that this second optimisation is not always successful — and if this occurs the outputs are returned as if the option was disabled. After processing was completed (*AddBiomechanics* kindly sends an e-mail to alert you), the output musculoskeletal models and results (i.e. inverse kinematics and dynamics — including the residual forces and moments about the pelvis) were downloaded from the application. Data were extracted for the same three gait cycles used in previous approaches and averaged across each participant for comparability. The computational time in the *AddBiomechanics* approach was calculated by reviewing the processing logs for each participant and summing the time in the two optimisations. Scaling of the computational time for the entire running trial was necessary for an accurate comparison to the other approaches where single gait cycles were processed. Therefore, the entire *AddBiomechanics* computational time was scaled by the relative duration of the entire running trial to the average duration of the three gait cycles for each participant.

## Results

### Computational Time

The mean (± standard deviation [SD]) computational times (in minutes) were 0.35 (± 0.08), 1.21 (± 0.20), 20.74 (± 4.14) and 1.37 (± 0.29) for RRA, RRA3, *MocoTrack* and *AddBiomechanics*, respectively (see Figure 1). The RRA and RRA3 approaches were the fastest, followed by *AddBiomechanics*, with *MocoTrack* taking approximately 4-60 times longer than all other approaches.

**Figure 1:**
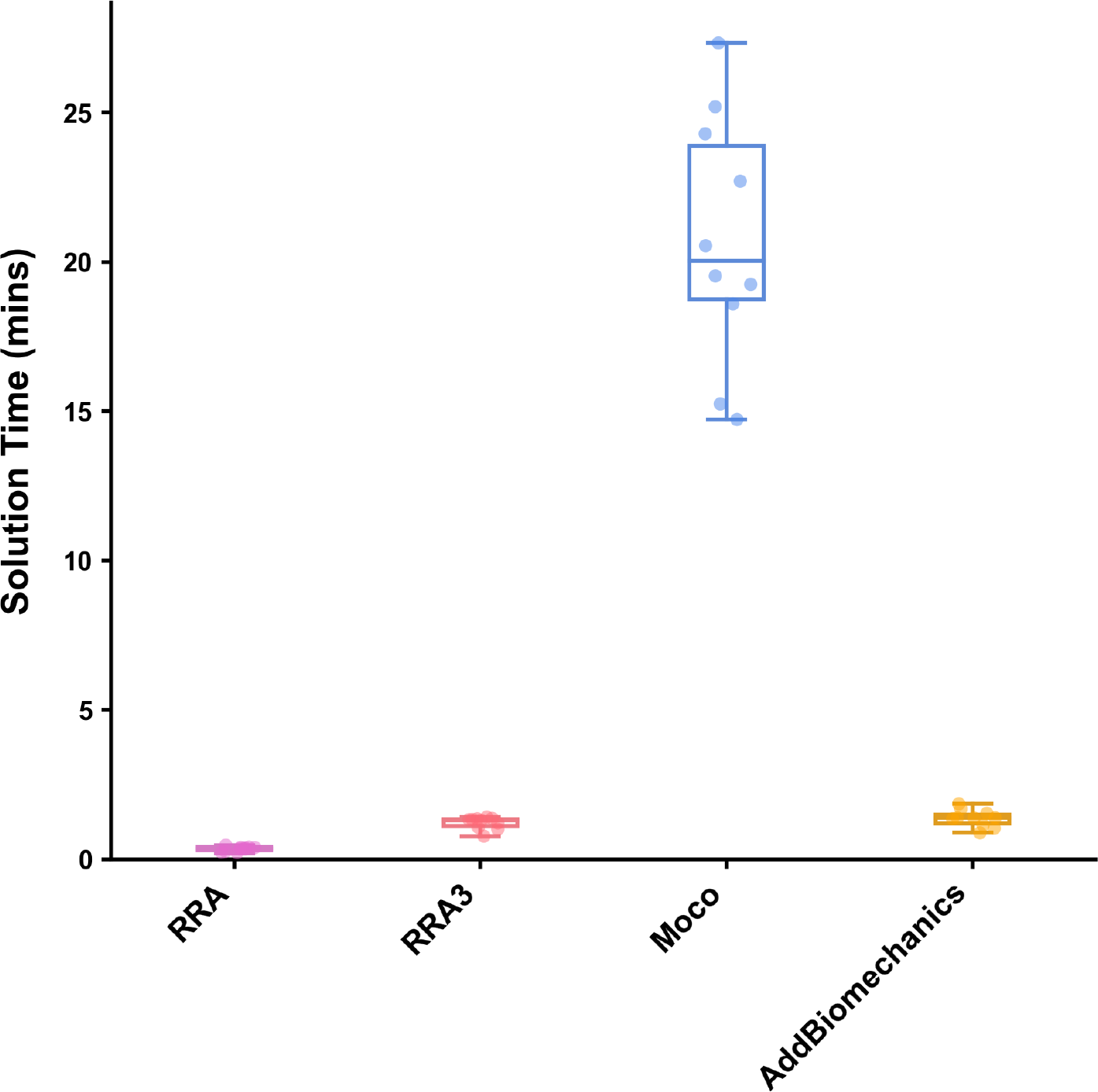
Solution times (in minutes) for processing a gait cycle using the Residual Reduction Algorithm (RRA — Purple), Iterative Residual Reduction Algorithm (RRA3 — Pink), MocoTracking (Moco — Blue), and AddBiomechanics (Gold) approaches. Horizontal lines within boxes equate to the median value, boxes indicate the 25^*th*^ to 75^*th*^ percentile, and whiskers indicate the range. Average solution times for each participants three gait cycles are displayed as points.

### Residual Forces

Average and peak residual forces (see Figure 2) were, on average, highest in the RRA approach (mean ± SD average residuals of 15.28 ± 3.20, 16.16 ± 5.52 and 13.96 ± 2.05 for FX, FY and FZ, respectively; mean ± SD peak residuals of 44.87 ± 6.63, 43.91 ± 12.05 and 43.55 ± 8.12 for FX, FY and FZ, respectively) followed by the RRA3 (average residuals of 9.15 ± 1.77, 8.96 ± 2.65 and 8.68 ± 1.58 for FX, FY and FZ, respectively; peak residuals of 26.78 ± 4.15, 27.63 ± 7.25 and 29.16 ± 8.18 for FX, FY and FZ, respectively) and *AddBiomechanics* (average residuals of 8.11 ± 21.35, 16.35 ± 42.53 and 7.33 ± 19.52 for FX, FY and FZ, respectively; peak residuals of 40.61 ± 76.45, 62.65 ± 111.19 and 25.39 ± 52.70 for FX, FY and FZ, respectively) approaches. In almost all cases, the *MocoTrack* approach recorded the lowest average and peak residual forces (average residuals of 0.14 ± 0.20, 0.33 ± 0.47 and 0.13 ± 0.13 for FX, FY and FZ, respectively; peak residuals of 0.32 ± 0.46, 0.66 ± 0.97 and 0.34 ± 0.37 for FX, FY and FZ, respectively). The RRA, RRA3 and *MocoTrack* approaches were able to achieve acceptable average and peak residual forces according to the threshold proposed by Hicks et al. [5] on average across all participants gait cycles. The *AddBiomechanics* approach also subceeded this threshold for the majority of participants, with the exception of one or two cases.

**Figure 2:**
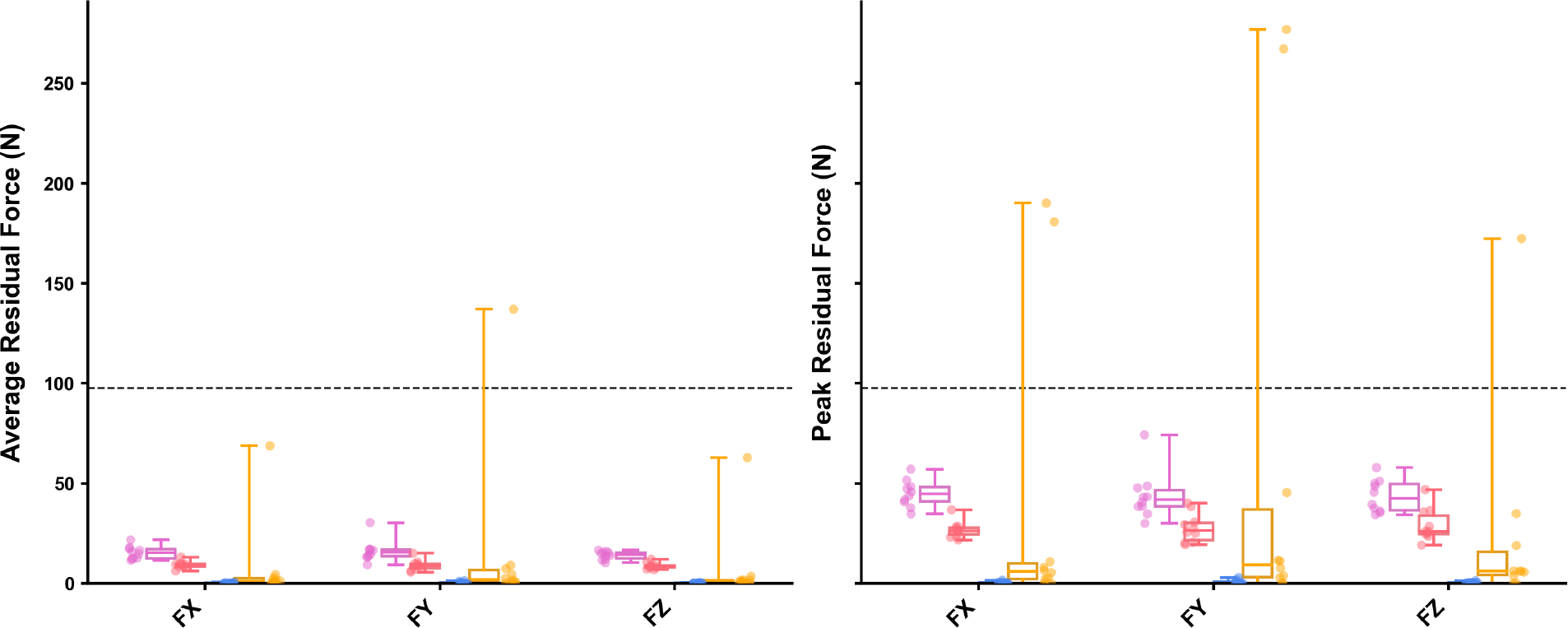
Average (left panel) and peak (right panel) residual forces (in Newtons [N]) for gait cycles processed using the Residual Reduction Algorithm (RRA — Purple), Iterative Residual Reduction Algorithm (RRA3 — Pink), MocoTracking (Moco — Blue), and AddBiomechanics (Gold) approaches. Horizontal lines within boxes equate to the median value, boxes indicate the 25*th* to 75^*th*^ percentile, and whiskers indicate the range. Average residual forces for each participants three gait cycles are displayed as points. The black dashed line represents the proposed acceptable threshold for residual forces.

### Residual Moments

Average and peak residual moments (see Figure 3) were typically highest in the RRA (mean ± SD average residuals of 26.72 ± 5.13, 24.57 ± 2.89 and 43.36 ± 7.90 for MX, MY and MZ, respectively; mean ± SD peak residuals of 83.02 ± 19.02, 70.84 ± 10.88 and 129.75 ± 20.66 for MX, MY and MZ, respectively) and *AddBiomechanics* (average residuals of 27.29 ± 6.01, 16.07 ± 4.91 and 27.84 ± 6.81 for MX, MY and MZ, respectively; peak residuals of 79.94 ± 15.63, 43.00 ± 11.69 and 90.56 ± 20.83 for MX, MY and MZ, respectively) approaches, followed by the RRA3 approach (average residuals of 12.71 ± 3.17, 18.86 ± 2.50 and 24.59 ± 4.90 for MX, MY and MZ, respectively; peak residuals of 44.77 ± 11.25, 57.96 ± 18.42 and 74.63 ± 12.12 for MX, MY and MZ, respectively). The *MocoTrack* approach recorded the lowest average and peak residual moments (average residuals of 0.33 ± 0.30, 0.85 ± 0.54 and 0.40 ± 0.41 for MX, MY and MZ, respectively; peak residuals of 0.90 ± 0.79, 2.30 ± 1.72 and 0.95 ± 0.91 for MX, MY and MZ, respectively) in all participants. Only the *MocoTrack* approach was able to consistently achieve acceptable average and peak residual moments according to the threshold proposed by Hicks et al. [5] across all participants gait cycles. Average residual moments from the RRA, RRA3 and *AddBiomechanics* approaches rarely subceeded, while the peak residual moments were always above this threshold

**Figure 3:**
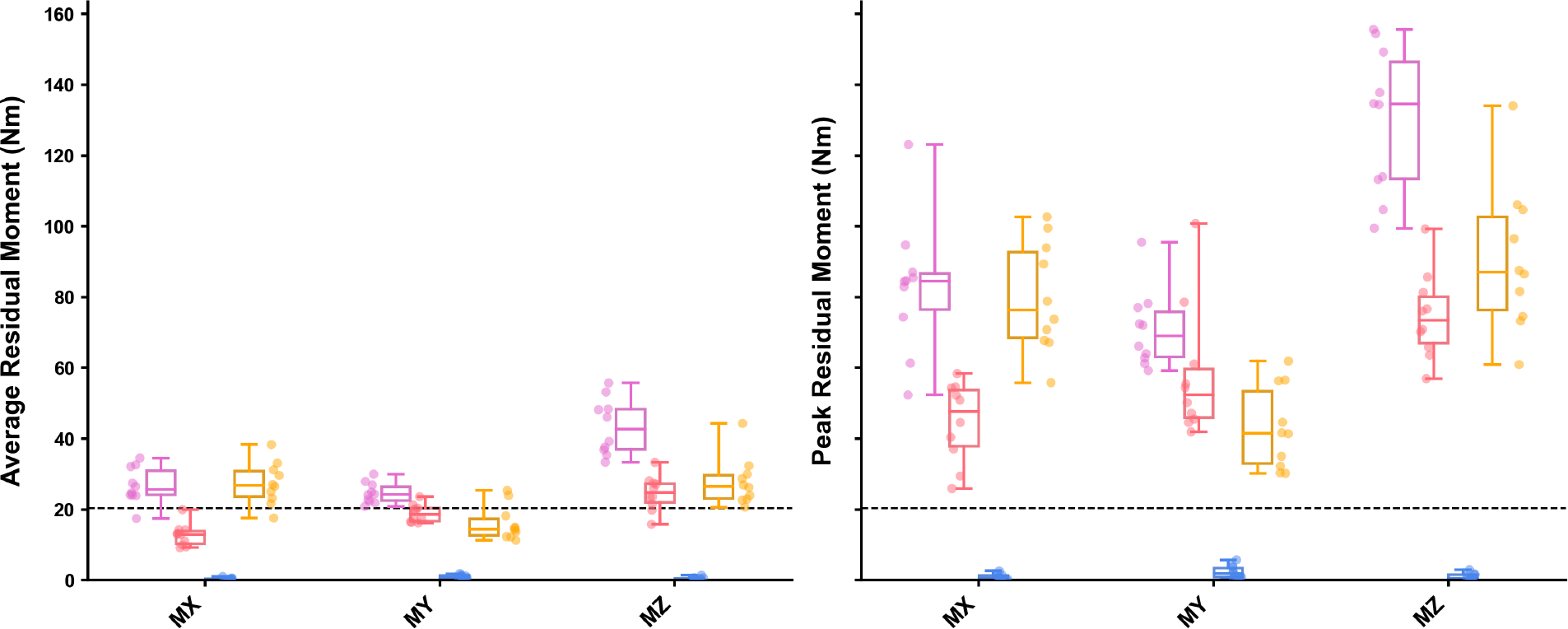
Average (left panel) and peak (right panel) residual moments (in Newton-Metres [Nm]) for gait cycles processed using the Residual Reduction Algorithm (RRA — Purple), Iterative Residual Reduction Algorithm (RRA3 — Pink), MocoTracking (Moco — Blue), and AddBiomechanics (Gold) approaches. Horizontal lines within boxes equate to the median value, boxes indicate the 25^*th*^ to 75^*th*^ percentile, and whiskers indicate the range. Average residual moments for each participants three gait cycles are displayed as points. The black dashed line represents the proposed acceptable threshold for residual moments.

### Joint Kinematics

Average joint kinematics were qualitatively similar across approaches for the majority of joint coordinates (see Figure 4). The greatest kinematic variations were for pelvic tilt, hip adduction/abduction and internal/external rotation, and ankle plantarflexion/dorsiflexion between *AddBiomechanics* and the other approaches; and for upper body kinematics (i.e. shoulder and elbow joint angles) between *MocoTrack* and the other approaches.

**Figure 4:**
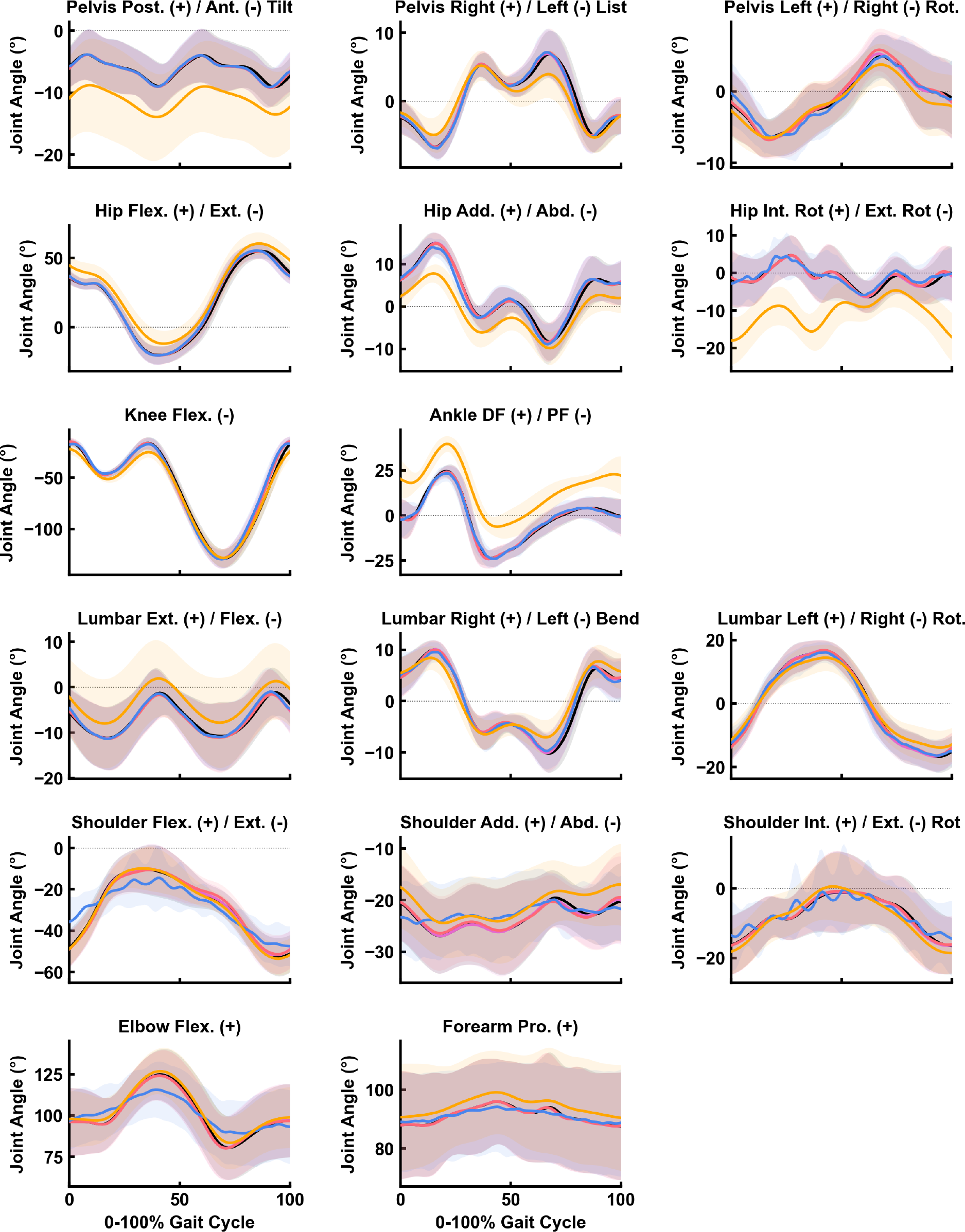
Mean (solid lines) ± standard deviation (shaded areas) of joint kinematics (in degrees) for gait cycles processed using Inverse Kinematics (IK), Residual Reduction Algorithm (RRA — Purple), Iterative Residual Reduction Algorithm (RRA3 — Pink), MocoTracking (Moco — Blue), and AddBiomechanics (Gold) approaches.

### Joint Kinetics

Average joint kinetics were qualitatively similar across approaches for all joint moments, with the exception of significant noise appearing in the *MocoTrack* signals (see Figure 5).

**Figure 5:**
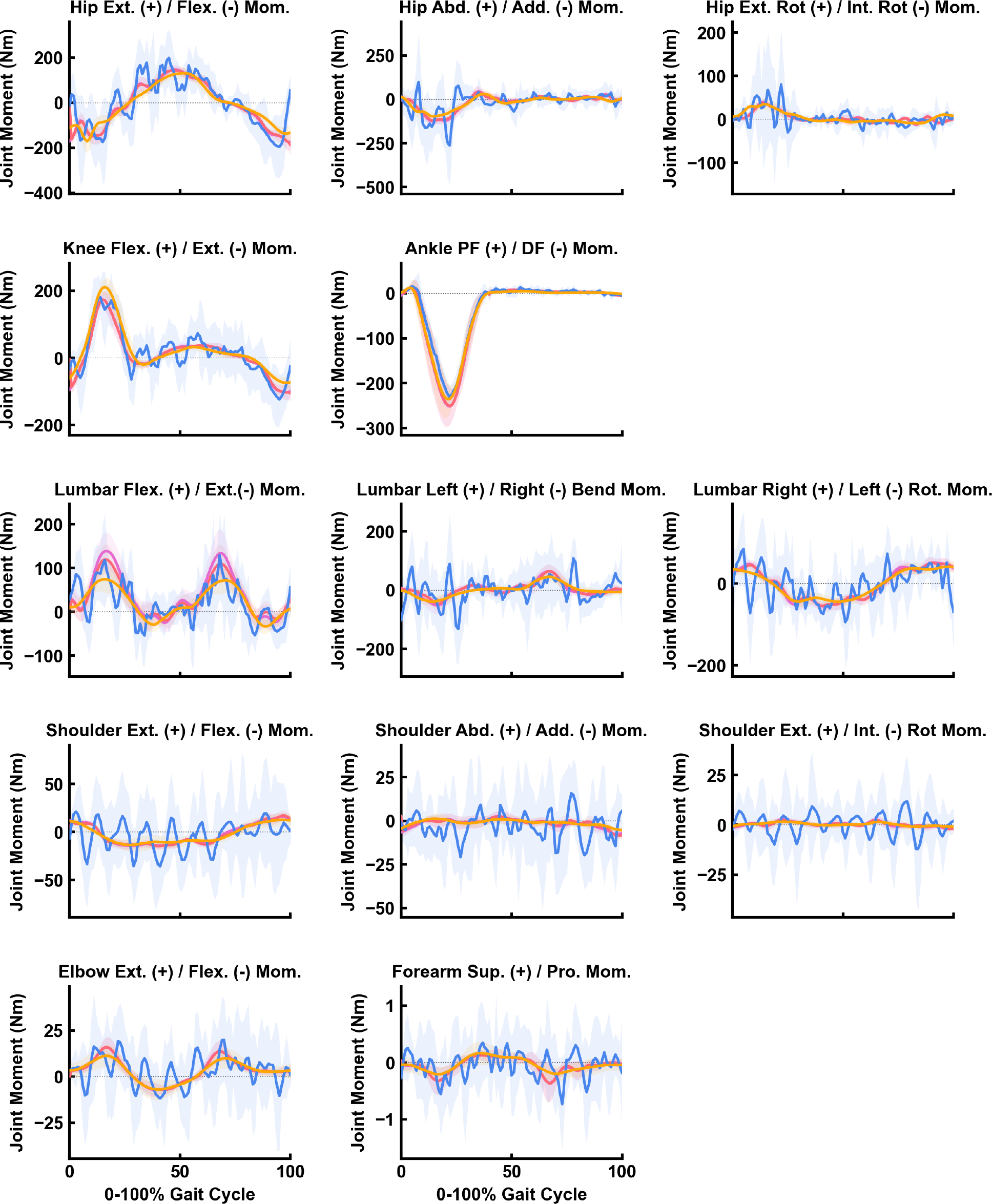
Mean (solid lines) ± standard deviation (shaded areas) of joint moments (in Newton-Metres [Nm]) for gait cycles processed using the Residual Reduction Algorithm (RRA — Purple), Iterative Residual Reduction Algorithm (RRA3 — Pink), MocoTracking (Moco — Blue), and AddBiomechanics (Gold) approaches.

## Discussion

Dynamic inconsistencies in simulations of human running can lead to unrealistic musculoskeletal model outputs (e.g. joint moments, muscle forces). Minimising the residual forces and moments at a models root segment is subsequently a recommended step within simulation pipelines [2, 5]. OpenSim — likely the most widely used musculoskeletal modelling and simulation software — now offers a variety of approaches applicable to residual reduction, yet these have never undergone a comprehensive comparison. This study aimed to compare the computational times, resultant residual forces and moments, and output joint kinematics and kinetics of different residual reduction approaches — or in more extravagant terms, set out on a quest for dynamic consistency in simulations of human running. A clear computational cost to residual reduction trade-off was identified, where approaches that required longer computational times were more effective at minimising residual forces and moments. In the majority of simulations, all approaches were able to reduce residual forces below recommended thresholds. However, only the *MocoTrack* approach was able to consistently achieve acceptable levels for residual moments. Minimal qualitative differences were observed in the resultant joint kinematics, with the exception of the *AddBiomechanics* and *MocoTrack* approaches producing variable lower and upper body kinematics, respectively, versus the remaining approaches. Joint kinetics were qualitatively similar between approaches despite the *MocoTrack* approach generating much noisier signals.

The present study revealed a clear trade-off between computational time and the capacity to reduce residual forces and moments. *MocoTrack* was by far the best and most consistent approach for reducing residuals, achieving near-zero levels (see Figures 2 and 3), and was the only approach that consistently reduced both residual forces and moments below recommended thresholds [5]. However, the *MocoTrack* approach took approximately 4-60 times longer than all others on average. Although *MocoTrack* had the highest computational times in the present study, the approximate 15-30 minute time-range substantially outperforms previous efforts [9, 11] to optimise the RRA process. It is likely that studies with a smaller sample (i.e. lower participant numbers and gait cycles to process) would be able to implement the *MocoTrack* approach, yet substantially larger studies may need to consider the longer computational times. The lower computational time of the *AddBiomechanics* approach must also be considered in the context of how it was assessed. An estimate of the *AddBiomechanics* processing time was taken from the processing logs in the application — and hence does not include the time taken to upload the data, how long the data was queued on the computing cluster, and the time taken to download the processed data. Uploads and downloads typically took less than a few minutes, while the cluster queue times were more variable and hence could add up for studies with large samples. Alternatively, users could consider these additional time costs as being offset against the unique time-saving aspects of using *AddBiomechanics* (e.g. outsourcing data processing, and removing the need for static trials and model scaling).

The joint kinematics produced by all approaches were, for the most part, qualitatively similar to one another (see Figure 4). The major exceptions were the pelvic tilt, frontal and transverse plane hip, and sagittal plane ankle angles from *AddBiomechanics* versus other approaches; and the shoulder and elbow joint angles from *MocoTrack* versus other approaches. While visualising the overall average motion demonstrates the relative whole-body consistency across approaches (see Figure 6), the specific differences at certain joints are also evident. The potential for variation in joint kinematics highlights the need to consider the residual reduction approach used when comparing results across studies, particularly for the joint angles that more largely varied. It is difficult to determine which approach achieved the most ‘accurate’ joint kinematics given the lack of a gold-standard measure to evaluate against. The inverse kinematic (IK) and *AddBiomechanics* joint kinematic solutions are derived by tracking experimental marker data, and therefore marker tracking error is a potential measure of accuracy for these approaches. However, marker error may not be a valid comparison to the RRA and *MocoTrack* approaches. The joint kinematics for RRA/RRA3 and *MocoTrack* were derived from tracking the IK results, and hence any initial marker errors in the IK data would likely propagate forwards and potentially increase in these solutions. Directly tracking marker data (instead of joint coordinates) alongside GRFs is a potential option within the *MocoTrack* framework [7]. Using a marker-tracking approach could also be valid in reducing residuals in running simulations and provide a more accurate comparison to the other marker-based approaches (i.e. IK and *AddBiomechanics*), however was not considered in the present study.

**Figure 6:**
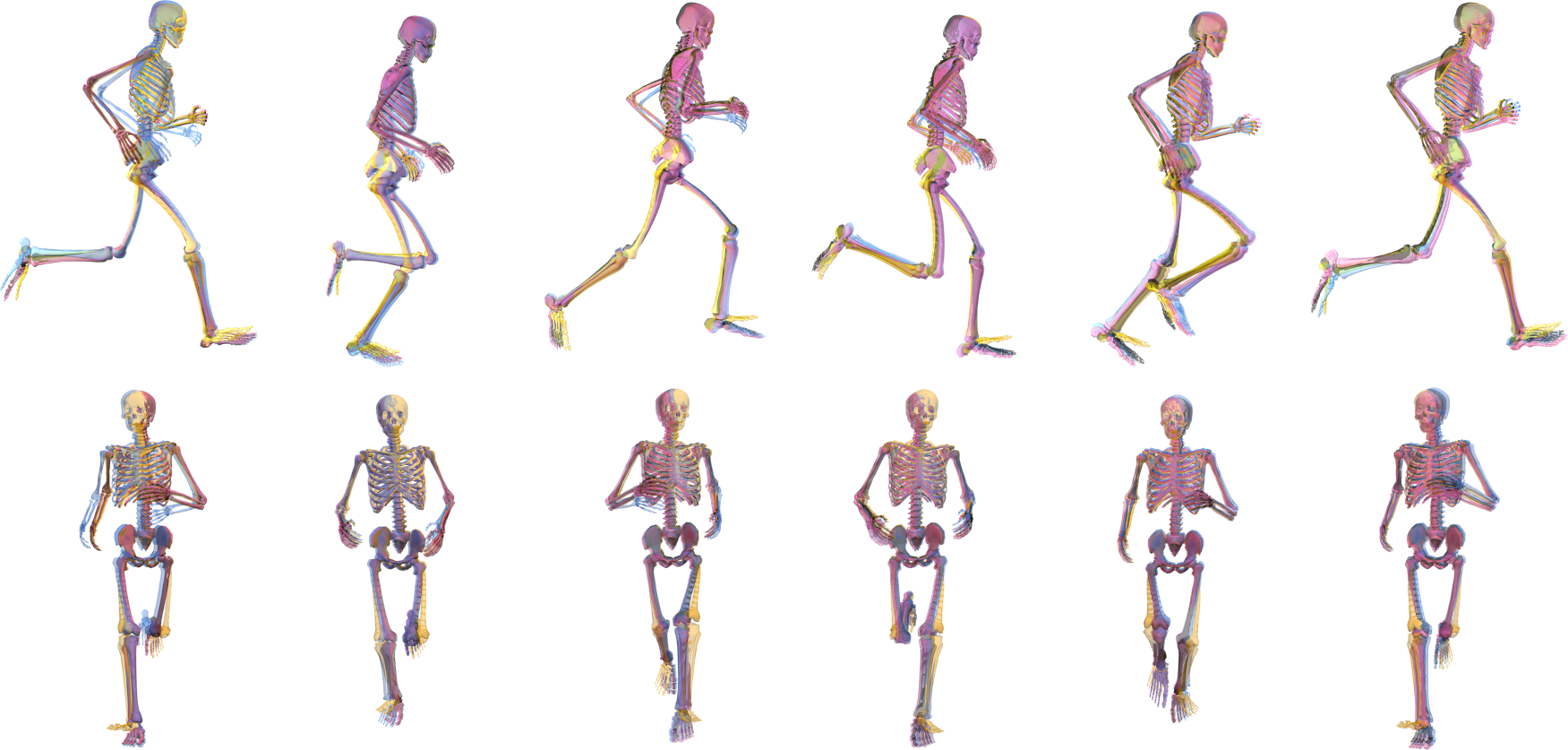
Group average joint motion at from the Residual Reduction Algorithm (RRA — Purple), Iterative Residual Reduction Algorithm (RRA3 — Pink), MocoTracking (Moco — Blue), and AddBiomechanics (Gold) approaches in the sagittal and frontal planes.

Similar to joint kinematics, the joint kinetics produced by all approaches were qualitatively similar (see Figure 5). However, the joint kinetics from *MocoTrack* were substantially noisier than those from all other approaches. Using a smaller time-step in torque-driven simulations can generate noisier signals from actuators driving joint motions, given the larger gap between sample points potentially requiring more abrupt shifts in the joint moments required. A smaller time-step was selected for the *MocoTrack* (i.e. 0.01s) versus RRA/RRA3 approaches (i.e. 0.0001s) in the present study for practical reasons, while the time-step in the *AddBiomechanics* approach is automatically determined based on the sampling rate of marker data (that being 0.01s in the dataset used). If the smaller time-step was a valid explanation for the noisier joint moments from *MocoTrack*, it could be assumed that the same phenomenon would present in the *AddBiomechanics* outputs — yet this was not the case. A more likely explanation is that the noise captured in the joint kinetics of the *MocoTrack* solution is simply being captured elsewhere in the other approaches. This begs the question, where did the noise go? The residual forces and moments that remained in the RRA, RRA3 and *AddBiomechanics* solutions were inherently noisy (see Figure 7 for an individual participant example), and perhaps offer a reason for why these approaches were able to achieve much smoother joint kinetic signals. The musculoskeletal models used in the present study were rigid in nature, which subsequently ignores soft tissue motion in the dynamics analyses and may offer an explanation for the lingering noise present in the simulations. Combining residual reduction approaches with more complex models (e.g. that include wobbling masses) may better capture joint dynamics while accounting for soft tissue motion [6].

**Figure 7:**
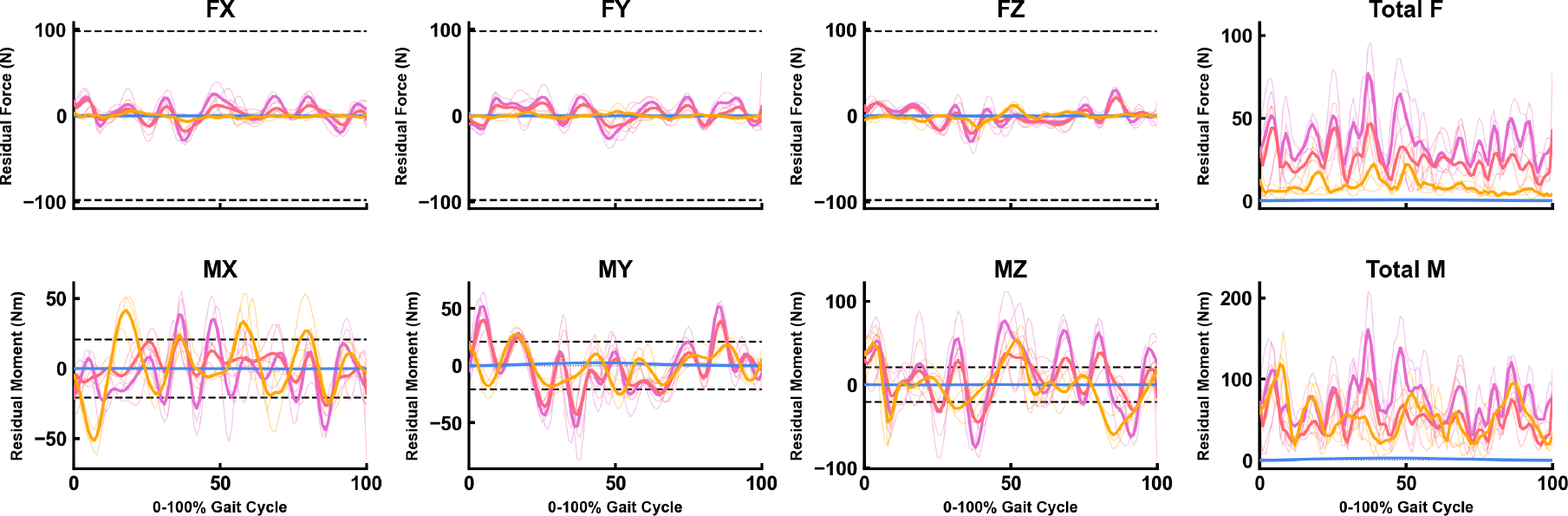
Average (thick line) and individual gait cycle (thin lines) residual forces (FX, FY, FZ and Total F) and moments (MX, MY, MZ and Total M) for the Residual Reduction Algorithm (RRA — Purple), Iterative Residual Reduction Algorithm (RRA3 — Pink), MocoTracking (Moco — Blue), and AddBiomechanics (Gold) approaches from a single participant example.

For the sake of complete honesty, very little effort went into selection of the input parameters and settings used in the various approaches. Where possible, the parameters used in the original work of Hamner and Delp [4] were replicated (i.e. tracking task weights and torque actuator settings in RRA) and reproduced in other approaches (i.e. RRA3 and *MocoTrack*). There are few objective recommendations for selecting these settings, hence the replication of the original studies [4] approach aimed to eliminate any additional subjectivity being added. Similarly, the default parameters in the *AddBiomechanics* application (e.g. the weight of residuals in the main optimisation) were not altered in any way. The results from the present study should therefore be considered with respect to the settings and parameters used for each approach. Past work has demonstrated that optimising the settings within residual reduction approaches can improve the dynamic consistency of the simulation, typically at the cost of additional computational time used to identify the optimised settings [9, 11]. Therefore, there may be room for improvement within the residual forces and moments achieved in the present study — yet users must consider the added computational time to deduce these. The time-frames and residuals reported in previous work optimising the RRA approach [9, 11] still exceed those of the best performing *MocoTrack* approach in the present study (i.e. ∼20 minutes for near-zero residuals with *MocoTrack*) — highlighting the effectiveness of this method even with minimal consideration of the input settings and parameters.

While the findings of the present study could support a broad recommendation for the *MocoTrack* approach, there is a practical user-based element to consider with respect to wide-spread adoption. At present, OpenSim’s Moco tools are only accessible via C++, MATLAB or Python scripting. In contrast, the RRA approaches can be accessed in this way plus via the graphical user interface (GUI); while the *AddBiomechanics* approach is accessible via a simple-to-use web application. Therefore, only OpenSim users with appropriate scripting knowledge and skills could theoretically implement the *MocoTrack* approach.

There is no information as to what proportion of the OpenSim user base would fit this description, but it is unlikely to be everyone. The analysis code (in Python) from the present study is publicly available (see the associated SimTK project page at https://simtk.org/projects/dynamic-quest) to support implementation of similar approaches. However, integration of the Moco suite of tools into the OpenSim GUI would likely support further use.

### Limitations

The findings from this study must be considered within the scope of the work. Only treadmill running at a single speed, in a healthy population and using a relatively small sample size (*n* = 10) was examined. The singular fastest speed in the dataset (i.e. 5.0 m·s^-1^) was chosen given the expected higher forces and accelerations having the potential to generate larger residual forces and moments. The various residual reduction approaches examined likely have a similar ability across different running speeds, however the results from the present study cannot confirm or refute this. It is possible that the magnitude of difference in residuals between the approaches could reduce at slower running speeds or in slower gait tasks (e.g. walking) — potentially making some of the lesser performing approaches more valid in these contexts. Similarly, there may be some variation in the success of the residual reduction approaches when examining overground instead of treadmill running. Given a single gait cycle was processed for most residual reduction approaches, it is expected that any difference between these running modalities would be minimal. The *AddBiomechanics* approach, however, may be the most affected given the whole-trial processing and potentially limited number of gait cycles that could be captured in a single overground trial. Lastly, performance of the residual reduction approaches may vary when examining altered gait in clinical populations (e.g. crouch gait or toe-walking in cerebral palsy).

## Conclusions

This study set out on a quest for dynamic consistency in simulations of human running by examining different residual reduction approaches (single and iterative RRA, *MocoTrack* and *AddBiomechanics*) available to OpenSim users. A computational time to residual reduction trade-off was identified, where approaches that took longer were more effective in reducing residual forces and moments. *MocoTrack* was the most consistent and best performing approach for reducing residuals to near-zero levels, however required substantially longer computational times and produced noisier joint kinetic signals. Joint kinematics were mostly similar across the residual reduction approaches, however specific joint angle variations occurred with certain approaches. The findings from the present study provide a comprehensive analysis of the simulation outputs when using different residual reduction approaches in OpenSim, providing users with evidence to inform decision-making at the residual reduction step of their modelling and simulation workflow when analysing human running.

## Acknowledgements

Thanks must go to those who asked questions and discussed this work at the 2023 International Symposium on Computer Simulation in Biomechanics (organised by the Technical Group on Computer Simulation of the International Society of Biomechanics) — with special thanks to Ajay Seth for proposing a new, more extravagant title.

